# High through-put identification of miR-145 targets in human articular chondrocytes

**DOI:** 10.1101/2020.02.03.931683

**Authors:** Aida Martinez-Sanchez, Stefano Lazzarano, Eshita Sharma, Chris L. Murphy

## Abstract

**Objective:** MicroRNAs play a key role in biological processes, including cartilage development and homeostasis and are dysregulated in many diseases, including osteoarthritis. MiR-145 modulation induces profound changes in the human articular chondrocyte (HAC) phenotype, partially through direct repression of *SOX9*. Since miRNAs can simultaneously silence multiple targets, we aimed to identify the whole targetome of miR-145 in HACs. This information is critical if miR-145 is to be considered a target for cartilage repair.

**Methods:** RIP-seq (RNA-immunoprecipitation plus HT-sequencing) of miRISC (miRNA-induced silencing complex) was performed in HACs overexpressing miR-145 to identify miR-145 direct targets. The motif discovery method cWords was used to assess enrichment on miR-145 seed matches in the identified targets. RT-qPCR, Western (immuno-)blot and luciferase assays were used to validate miRNA-target interactions.

**Results:** MiR-145 overexpression affects the expression of over 350 genes and directly targets more than 50 mRNAs through the 3’UTR or, more commonly, the coding region.

We also demonstrate that miR-145 targets DUSP6, involved in cartilage organization and development, at the translational level. DUSP6 depletion using specific siRNAs lead to MMP13 up-regulation, suggesting that miR-145-mediated DUSP6 depletion contributes to the effect of miR-145 on MMP13 expression.

**Conclusion:** We demonstrate that miR-145 directly targets several genes in primary chondrocytes including those involved in the expression of the extracellular matrix and inflammation. Thus, we propose miR-145 as an important regulator of chondrocyte function and a new target for cartilage repair.

---

The articular chondrocyte is the only cell type in the articular cartilage, solely responsible of synthetizing the components of the extracellular matrix (ECM) that allows an almost friction-free painless movement of the joint. When the articular chondrocyte function is altered, the structure of the ECM can be compromised leading to the development osteoarthritis (OA). Currently, there is no cure for OA, which affects over 250 million people worldwide (WHO http://bjdonline.org/), and most common treatments can only aim to pain reduction. The normal healthy articular chondrocyte phenotype maintains the equilibrium between anabolic molecules, such as Collagen type II (COL2A1) - a main component of the ECM- and catabolic enzymes such as collagen-degrading metalloproteinase MMP13 (1, 2). Understanding the mechanisms underlying human articular chondrocyte function (HAC) is indispensable to develop therapies aiming to restore the capacity of the HAC to produce a balanced ECM.

MicroRNAs (miRNAs) are small RNAs that silence gene expression post-transcriptionally (3). Over 2000 miRNAs are expressed in humans (4) that play regulatory roles in most biological processes, development and disease (5–7). MiRNAs have been widely shown to not only regulate cartilage development but also modulate the normal function of the adult articular chondrocyte (8–10). We have previously identified miR-145 to profoundly impact the expression of essential components of the ECM, such as ACAN, COL2A1 or MMP13 at least partially by directly targeting the master regulator, transcription factor SOX9 (11). Interestingly, miR-145 expression is up-regulated in OA chondrocytes (12) raising the possibility of inhibiting miR-145 as a promising approach for cartilage repair. Nevertheless, a single miRNA typically targets dozens to even hundreds of mRNAs simultaneously (3), which could result in unpredicted phenotypes upon manipulation. Therefore, if miR-145 is going to be considered as a target for cartilage repair, it is fundamental to determine its whole targetome in HACs. Current available computational methods for miRNA target determination rely in the extent of the complementarity between the seed region of the miRNA (6 nucleotides positions 2±7) and the 3’UTR of the mRNAs, which results in hundreds of predictions, most of which don’t represent bona-fide interactions. Additionally, efficiency of these tools tends to be poor for the detection of non-conserved binding sites, binding sites with poor pairing or for those located outside the 3’UTR of the mRNAs. Moreover, computational approaches don’t take into account the possibility of tissue-specific interactions (13, 14).

Therefore we aimed to determine miR-145 targets in HACs using an un-biased high-throughput experimental approach capable of detecting actual miRNA-mRNA interactions(14). Accordingly, in the present study we performed RNA/miRISC-immunoprecipitation experiments (15–17) followed by high-throughput sequencing (RIP-seq) in primary HACs isolated from healthy human cartilage and previously transfected with a miR-145 mimic. Using this method, we detected 59 mRNAs enriched in miRISC immunoprecipitates upon miR-145 over-expression, indicating direct miRNA-mRNA interaction. Reporter assays confirmed that miR-145 silences the expression of several of these genes through sequences present in their 3’UTR or coding sequence. One such mRNA identified, DUSP6, had previously been reported to be involved in cartilage function. The predicted miR-145 binding site in DUSP6 was validated by reporter assays and, as expected, DUSP6 protein levels were modulated in primary HACs upon miR-145 over-expression/inhibition. Both miR-145- and siRNA-mediated downregulation of DUSP6 resulted in increased expression of the catabolic gene MMP13, indicating that DUSP6 is at least partially responsible of the effect that miR-145 exerts in MMP13 gene expression.

## MATERIALS AND METHODS

### Cell culture and transfection

Following local ethics committee guidelines and after informed consent, healthy human articular cartilage was extracted and HACs were isolated and cultured as previously described (11). Primary cells (passage 0 - P0) were harvested after 5-7 days of culture or sub-cultured for a further 5-7 days and passaged one, two or three times (P1-P3). HACs were transfected with control or hsa-miR-145 miRCURY LNA Power inhibitors (Exiqon) or control or DUSP6 siRNA (ThermoScientific) as previously described (18).

HeLa cells were transfected using 100ng of plasmids together with 5nM control or miR-145 mimics. After 24 hours, cells were lysed and luciferase activity determined with the Dual-Glo luciferase assay system (Promega).

### Plasmids

The sequence containing three consecutive perfect matches for hsa-miR-145-5p was excised from pSG5-Luc-3xmiR-145 (11) and sub-cloned in pMirGlo (Promega) to generate pMiRGlo 3xmiR-145.

MOK 3’UTR, MOK CDS, CYR61 3’UTR, HSBP1 3’UTR, RTKN 3’UTR, CTBP1 3’UTR, CTBP1 CDS, FKBP2 CDS, HIST4E full-length, GLT8D1 3’UTR, RAB7A 3’UTR, RAB7A CDS, PLOD1 3’UTR, SEC11A 3’UTR, FN3K CDS, UXS1 3’UTR and DUSP6 3’UTR were amplified by PCR using specific oligonucleotides (sequences available upon request) from HACS cDNA, inserted into the pGEM-T vector (Promega) and subcloned into pMiRGlo.

Three point mutations in the predicted miR-145 seed binding site in DUSP6 were introduced in pGEM-T-DUSP6 3’UTR using a Phusion^®^ site-directed mutagenesis kit (NEB) and the mutagenic primers: Forward: 5’-TGTGAGCATGGGTACCCATT-3’ and Reverse 5’- CACACACACTTCGTCTTTTATACAAA-3’. Mutated DUSP6 3’UTR was sub-cloned into pMiRGlo. All the constructs were verified by sequencing.

### Ribonucleoprotein Immunoprecipitation, mRNA/miRNA isolation and RT-qPCR

Ribonucleoprotein Immunoprecipitation was performed as described previously (17) from 30×10^6^ P3 chondrocytes from three different patients (19yo., female; 8yo., male; 28yo., male) transfected with miR-145 or control mimics for 24h using 90 μl Sepharose G beads (Abcam, Cambridge, UK) pre-coated with 6ug of mouse anti-human Ago2 (clone2E12-1C9, Abnova, Taipei City, Taiwan) or mouse IgG isotype control (Abcam). RNA was eluted from the beads and was isolated using TRIzol. Total RNA was extracted using TRIzol from a 10% volume of the samples submitted to immunoprecipitation, prior incubation with Ago2 antibody.

For miRNA detection, 5% of the recovered RNA was reverse transcribed using the TaqMan miRNA RT kit (Life Technologies) and miR-145, miR-140 and RNU24-specific primers (Life Techonologies). QPCR was performed with a TaqMan master mix and the appropriate TaqMan probes (Life Technologies). The remaining RNA was treated with DNAseI (Invitrogen) and sent for High-throughput sequencing.

Total RNA was extracted from HACs using TRIzol according to the manufacturer’s instructions. 200-500 ng of RNA were reverse transcribed using the High Capacity cDNA RT kit (Life Techonologies). QPCR followed, using a SYBR Green PCR master mix (Life Techonologies) and specific primers (sequences available upon request).

### mRNA-seq library construction, sequencing and analysis

RNA quantity and integrity were assessed using Quant-IT RiboGreen Kit (Invitrogen) and Agilent Tapestation 2200R6K. mRNA enrichment was achieved from 400-900ng of total RNA using a Magnetic mRNA Isolation Kit (NEB). Generation of double stranded cDNA and library construction were performed using NEBNext^®^ mRNA Sample Prep Reagent Set 1 (NEB). Ligation of adapters was performed using Adapters prepared at the WTCHG according to the Illumina design (Multiplexing Sample Preparation Oliogonucleotide Kit). Each library was subsequently size selected with two Ampure Bead bindings. The following custom primers were used for PCR enrichment: MultiplexPCRprimer1.0 5’-AATGATACGGCGACCACCGAGATCTACACTCTTTCCCTACACGACGCTCTTCCGA TCT-3’. Indexprimer: 5’-CAAGCAGAAGACGGCATACGAGAT[INDEX]CAGTGACTGGAGTTCAGACGTGTGCTCTTCCGATCT-3’. Indices were according to the eight bases tags developed by WTCHG(19). The following modifications to the described workflow were applied to RNA recovered after Ribonucleoprotein Immunoprecipitation. Firstly, no polyA selection or ribodepletion was applied. Secondly, the number of PCR cycles was increased from 12 to 15. Amplified libraries were analysed for size distribution using the Agilent Tapestation 2200 D1K. Libraries were quantified by quantitative RT-PCR using Agilent qPCR Library Quantification Kit and a MX3005P instrument (Agilent) and relative volumes were pooled accordingly. Finally, a second RT-qPCR was performed to measure the relative concentration of the pool compared to a previously sequenced mRNA library in order to determine the volume to use for sequencing. Sequencing was performed as 50bp paired end read on a HiSeq2000 according to Illumina specifications.

FASTQ files were generated for each sample and initial data quality checks of the raw sequence data were performed. Reads were then mapped to the human genome (GRCh37) using TopHat(20). HT-Seq(21) was used to summarise the mapped reads into a gene count table, the final version of which contained one column per sample. Ensembl (v65) annotations were used to define gene models. All subsequent data analysis was performed in R using relevant BioConductor packages(22).

Total RNA profiles were consistent and generated ~20 million reads mapping to Ensembl genes per sample. IP samples generated 2-3 million reads per sample and one potential outlier sample was observed on clustering and PCA plots. The R package ‘DESeq’(23) was used to normalise the samples for library depth and perform a paired analysis to find differentially expressed genes between the samples over-expressing miR-145 and controls. Total RNA and IP samples were treated as separate datasets. The two resulting gene lists were compared to identify genes that were significantly up-regulated in the IP samples while being significantly down-regulated or unchanged (not significant) in the total RNA samples as potential targets of miR-145. For the analysis performed with IP samples from only two patients, we investigated the effect of the outlier and the final IP list was generated excluding the patient with the outlier sample. Genes were filtered by expression levels and those with log2CPM>2 from total RNA and IP samples were independently ranked by fold change of expression and subjected to motif discovery method cWords in two separate analysis per list, one for enrichment in the 3’UTRs and the other in the CDS (24).

### Western blotting

HACs were lysed in RIPA Buffer and 10-30μg total protein extract subjected to SDS-polyacrilamide-gel electrophoresis and transferred to PVDF membranes. Monoclonal anti-Ago2 (Abnova), monoclonal anti-DUSP6 (Abcam), monoclonal anti-phospho-ERK (Sigma-Aldrich), polyclonal anti-ERK (SantaCruz Biotechnologies) and monoclonal anti-Tubulin (Sigma-Aldrich) antibodies were used.

### Statistical analysis

GraphPad Prism 6.0. was used for statistical analysis. Statistical significance was evaluated by two tailed paired student’s *t* test. All data are shown as mean ±standard error of the mean (S.E.M.). *P* values<0.05 were considered statistically significant.

## RESULTS

In order to determine miR-145 direct targets in human chondrocytes we performed mRNA/miRISC-immunoprecipitation followed by high-throughput sequencing (RIP-seq) with an anti-Ago2 antibody and a lysate from P3 HACs previously transfected with a miR-145 mimic or a control. The specificity of Ago2-immunoprecipitation was verified by Western blotting (Supplemental Figure 1A) and hundreds of miRNAs were detected in Ago2 immunoprecipitates in comparison with an IgG control (data not-shown). The amount of miR-145 associated with Ago2 was 40 fold higher in cells over-expressing miR-145 (Supplemental Figure 1B). Immunoprecipitated RNA (IP-RNA) was obtained from three independent experiments with chondrocytes isolated from different donors and prepared for high-throughput sequencing. To assess the overall impact of miR-145 over-expression in gene expression in HACs, we sequenced total RNA (T-RNA) preceding IP.

The expression of 352 genes was significantly changed (FDR<0.05) following miR-145 overexpression (T-RNA, Supplemental Table 1). Out of these genes, 80 were down-regulated (DT-RNA, Table 1) suggesting that most of the effects of miR-145 on gene expression are indirect. 96 genes were enriched in IP-RNA from samples over-expressing miR-145. Out of these genes, 59 (Table 2) were reduced or unchanged in the corresponding T-RNA samples. Since miRNAs normally act by decreasing target-mRNA stability or translation efficiency we considered those 59 mRNAs as miR-145 direct targets (D-IP). Although miRNAs can also mediate repression through “seedless” sites (25) and/or through the CDS (26), miR-145 direct targets are generally expected to contain miR-145 seed matches in the 3’UTRs. TargetScan (TargetScan (v6.2) (27) http://www.targetscan.org/) detected miR-145 target sites in 17 out of our 59 candidates (Table 2) whereas RNAhybrid, a bioinformatic tool that gives little weight to the seed region and permits the analysis of any given mRNA sequence, including CDS or 3’UTRs, identified miR-145 binding sites within the 3’UTR, the CDS and both the CDS and 3’UTR of 20, 25 and 12 candidate genes, respectively (Table 2). Finally, we used cWords (24) to analyse words enriched in 3’UTRs and CDS of IP-RNA (Supplemental Figure 2A). As expected, the word with strongest enrichment in CDSs of up-regulated genes corresponds to the 6mer-seed site of miR-145 (ACTGGA-Supplemental Figure 2B) and several shifted variants of the seed site were also enriched (28). Surprisingly, words corresponding to miR-145 seeds were not significantly enriched in 3’UTRs (Supplemental Figure 2C/D). Altogether, our data supports that those genes up-regulated in IP-RNA (D-IP) are miR-145 direct targets and that miR-145-target interactions occur predominantly through their CDS.

**Table 1.**
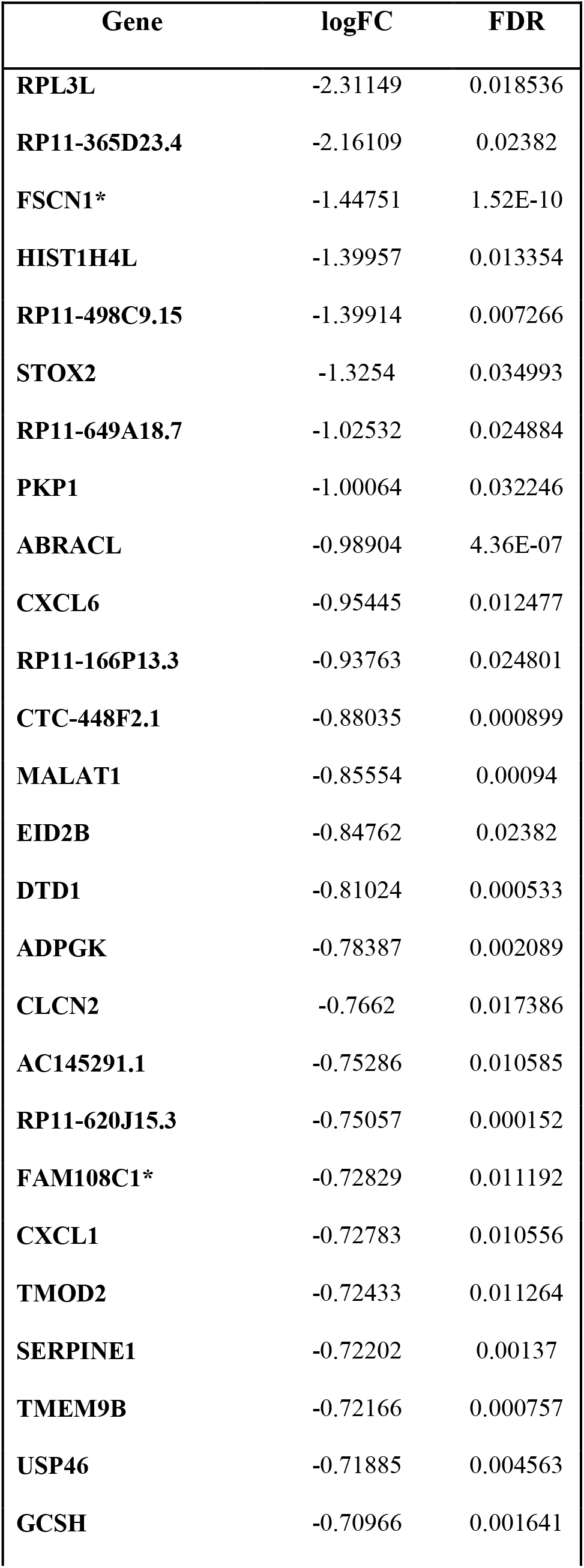
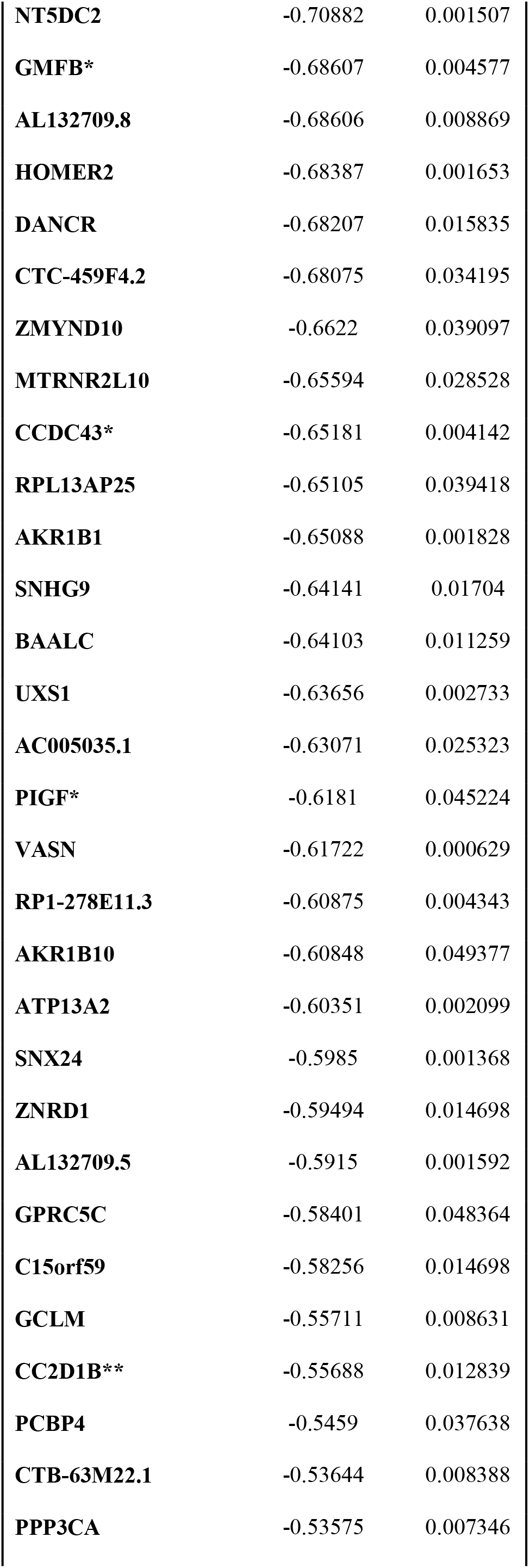
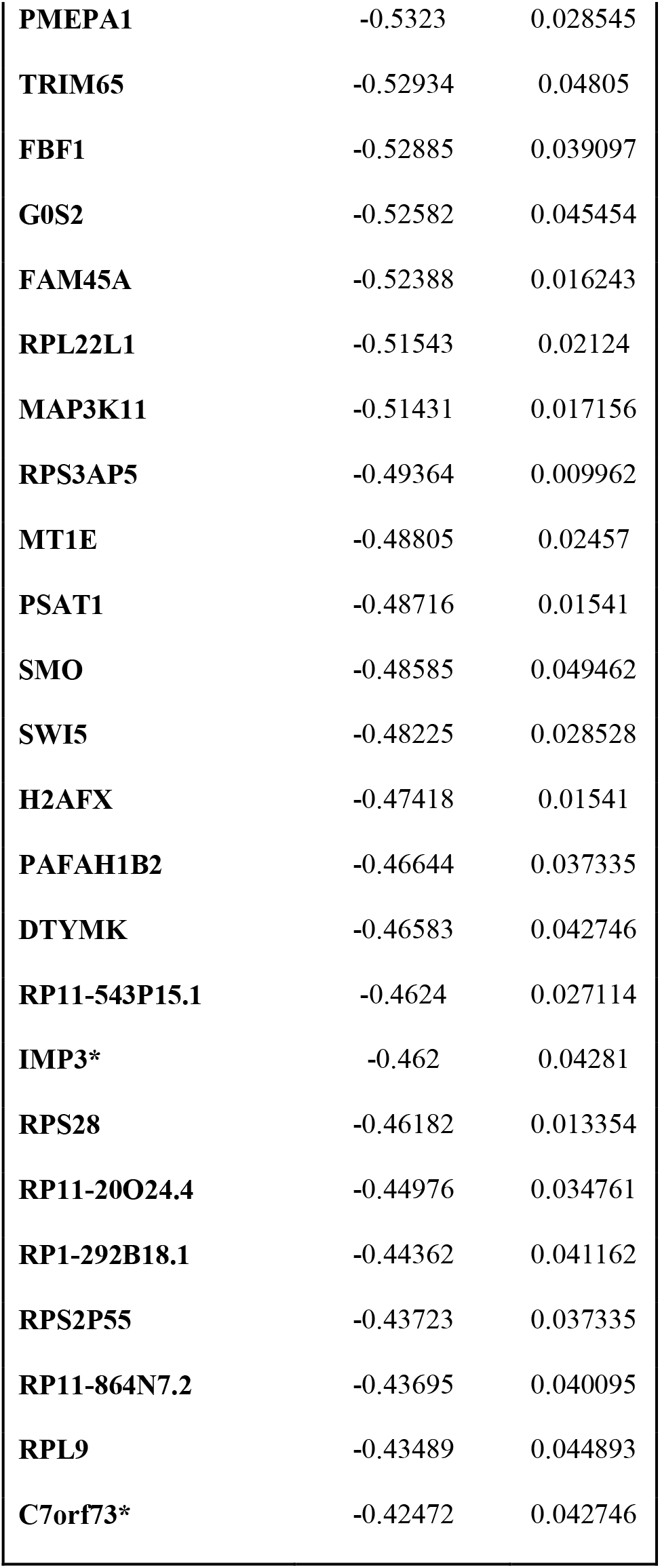
mRNAs significantly down-regulated (FDR<0.05) upon miR-145 overexpression. (TD-RNA: Total down-regulated mRNA). Control or miR-145 mimics were transfected in P3 HACs from three different patients. 24h after transfection, cells were lysed and RNA was extracted and submitted to HT-seq. **: genes significantly enriched in DT-IP; *: genes significantly enriched in Ago2-IP following miR-145 over-expression when the outlier patient was excluded from the analysis (Supp Table 2).

Further supporting that most of the changes occurred in gene expression upon miR-145 modulation are indirect, T-RNA genes were not enriched in miR-145 seeds. Out of the 80 genes which expression was down-regulated by the action of miR-145 (DT-RNA), only one was included in the D-IP group. Due to the complexity of the Immunoprecipitation protocol and the low amount of mRNA recovered, we couldn’t rule out the possibility that miR-145-mRNA weak interactions were lost. Indeed, clustering and PCA plots suggested one potential outlier. Thus, we expanded the analysis, excluding the outlier and generating additional lists of candidates, with genes that were significantly enriched in IP-RNA from only two patients and simultaneously reduced or unchanged in the corresponding T-RNA samples (Supplemental table 2). This additional list contained several genes which expression was repressed by miR-145 action (DT-RNA) (Table 1), and the gene Sox9, which we and others have previously proven to be a main direct target of miR-145 in HACs. Consistently, words corresponding to miR-145 seed were clearly enriched in both CDSs and 3’UTRs of these genes (Supplemental Figure 3).

HACs become gradually de-differentiated with passage and we therefore decided to validate our findings in freshly isolated (fully differentiated) HACs. We over-expressed miR-145 or a control mimic in P0 HACs and analysed the expression of several randomly-selected miR-145 direct targets by qPCR (Figure 1A). We selected 17 genes from the D-IP list, of which only one had been detected as down-regulated by miR-145 over-expression (DT-RNA). Nevertheless, when these candidates were examined in a higher number of patients and 48h (instead 24h), an additional 4 genes were significantly (p<0.5) down-regulated and 3 showed a tendency to be down-regulated (p<0.1), while only over half were consistently unchanged, suggesting that miR-145-target interaction results first in translational repression that is later on followed by mRNA destabilization.

**Figure 1.**
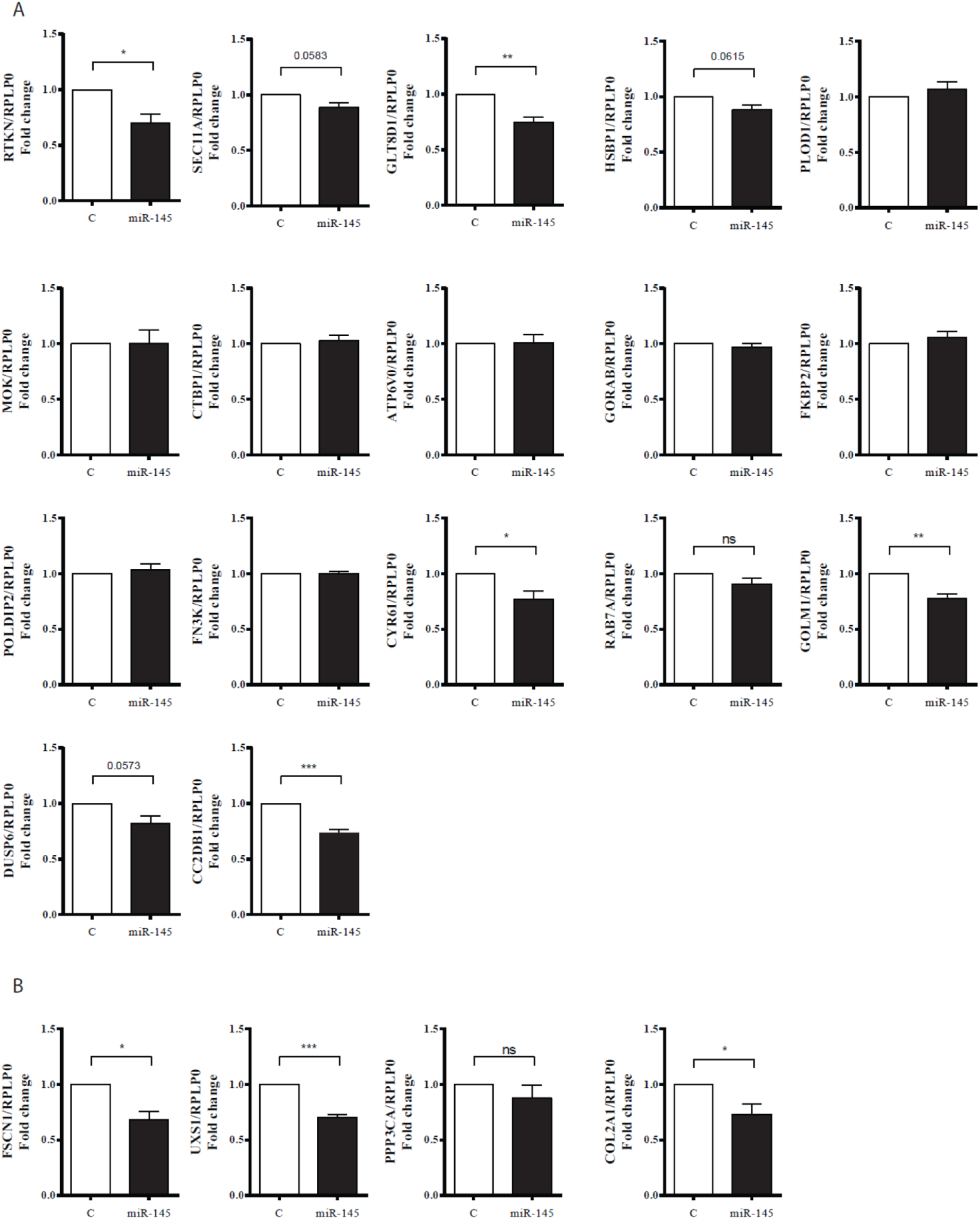
Analysis of the effect of miR-145 over-expression on mRNA levels of selected genes in freshly isolated HACs. **A)** 16 randomly selected mRNAs from CD-IP and CD2DB1 from both CD-IP and DT-RNA. **B)** 3 genes only present in DT-RNA and COL2A1. Control (C) or miR-145 mimics were transfected in freshly isolated HACs and RNA was extracted 48h after transfection. Values are presented as relative to that obtained in cells transfected with control mimics for each patient and normalized to RPLP0. Data represents average ± S.E.M. from 7 different experiments, each performed with different donor cells. \**P* < 0.05; \*\**P* < 0.01; \*\*\**P* < 0.001; ns, not significant.

We also validated the effect of miR-145 in three of the genes present in DT-RNA but not in D-IP, showing a significant effect for FSCN1 and UXS1, but not PPP3CA (Figure 1B). We additionally corroborated miR-145 effect in COL2A1, which we have previously shown to be indirectly regulated by miR-145 (Figure 1B).

To confirm an effect of miR-145 at the protein level, we generated luciferase reporters were the 3’UTR or CDS (or both) of 12 randomly selected D-IP mRNAs were cloned downstream the firefly luciferase ORF (Figure 2). Consistently, MOK, RAB7A and FN3K’s CDS, HSBP1 and PLOD1’s 3’UTRs, HIST4E’s full-length sequence and SEC11A’s both 3’UTR and CDS significantly mediated firefly luciferase silencing upon miR-145 over-expression. Only two out of the 12 sequences assayed (CTBP1 and FKBP2) failed to mediate silencing. Interestingly, no target sites were predicted for HIST4E mRNA, but miR-145 over-expression significantly reduced luciferase expression when the full-length HIST4E cDNA was cloned downstream its ORF, demonstrating that HIST4E is targeted by miR-145 and proving that our approach can detect targets that wouldn’t be predicted by even very permissive bioinformatics tools. Similar experiments were performed for CYR61, RTKN and GLT8D1 (which mRNA levels were down-regulated when assayed 48h after transfection, but not 24 (Figure 1A)) and we confirmed that their 3’UTRs mediated luciferase silencing, as expected.

**Figure 2.**
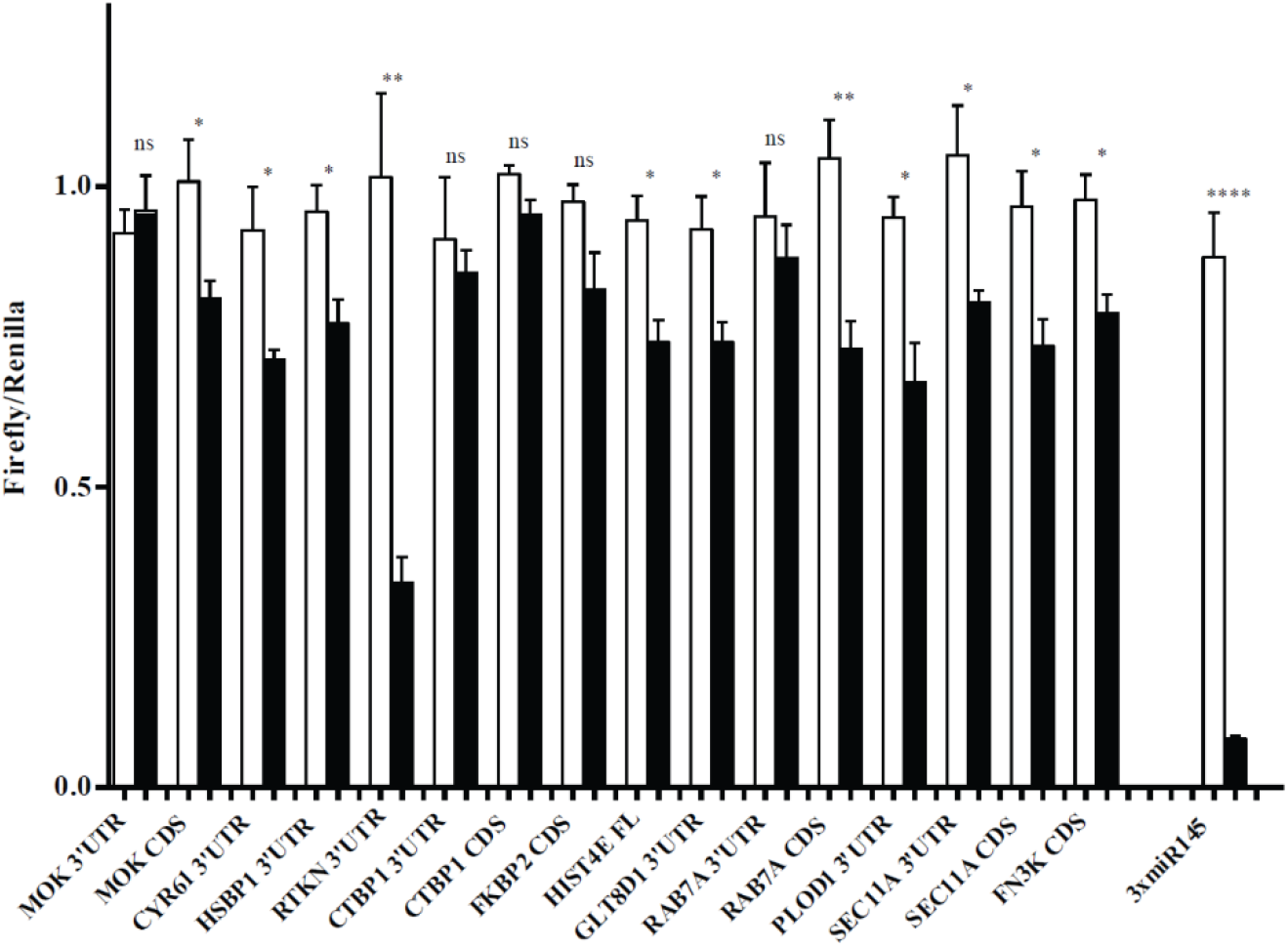
Luciferase reporter assays to validate miR-145 direct effect on the expression of selected genes. Hela cells were transfected with luciferase reporters containing full-length CDSs or 3’UTRs, or both, of 12 randomly selected mRNAs from CD-IP, as indicated, downstream the firefly luciferase ORF. Control (white bars) or miR-145 (black bars) mimics were co-transfected in the cells. Values are normalized to the levels of renilla luciferase, independently expressed by the same vector and are shown as relative to that obtained for each construct co-transfected with the control miRNA mimic (±S.E.M). As positive control, a construct containing three perfectly complementary binding sites for miR-145 (3xmiR-145) was included in the experiment. \**P* < 0.05; \*\*\*\**P* < 0.0001; ns, not significant.

In order to identify the biological processes and pathways in which miR-145 is involved, we submitted the full list of genes altered upon miR-145 overexpression (Supplemental Table 1, T-RNA) together with those included in the D-IP list (Table 2), to the Database for Annotation, visualization and Integrated Discovery (DAVID v 6.7). Significantly enriched pathways included immune response and proteolytic processes, both characteristic of cartilage disease (Supplemental Table 3) (p<0.1). One of the genes, DUSP6, encodes a dual-specificity phosphatase, MAP kinase phosphatase 3 (MKP3) that inhibits MAPK activity by dephosphorylating threonine and serine residues. MKP3 may regulate the elongation of cartilage elements of digit 2 during development through repression of FGF signalling (29). Interestingly, loss of DUSP6 in mouse embryos leads to dominant, incompletely penetrant and variable phenotypes including skeletal dwarfism (a common feature with Sox9 depletion), hearing loss and coronal craniosynostosis. In addition, chondrocytes in the mutant’s growth plate-proliferating zone were disorganized (30). Thus, we hypothesized that miR-145-mediated DUSP6 regulation affects adult chondrocyte function. As expected, miR-145 overexpression in HACs led to a strong reduction of DUSP6 protein (Figure 3A/B) in both normoxia or hypoxia, which promotes the differentiated phenotype. Conversely, prevention of miR-145 function with specific inhibitors resulted in increased DUSP6 expression (Figure 3C). As previously showed, miR-145 over-expression causes only a small, non-significant, decrease in DUSP6 mRNA (Figure 1A), suggesting that its effect occurs mainly at the translational level. A miR-145 binding site was predicted in the position 1018-1025 of DUSP6 3’UTR. In order to validate its functionality, we cloned DUSP6 3’UTR downstream the luciferase ORF. Transfection of miR-145 mimics significantly reduced luciferase activity of the wild-type construct but not that containing a 3-nucleotide point mutation in the miR-145-predicted target site (Figure 3D), validating the functionality of this predicted binding site.

**Figure 3.**
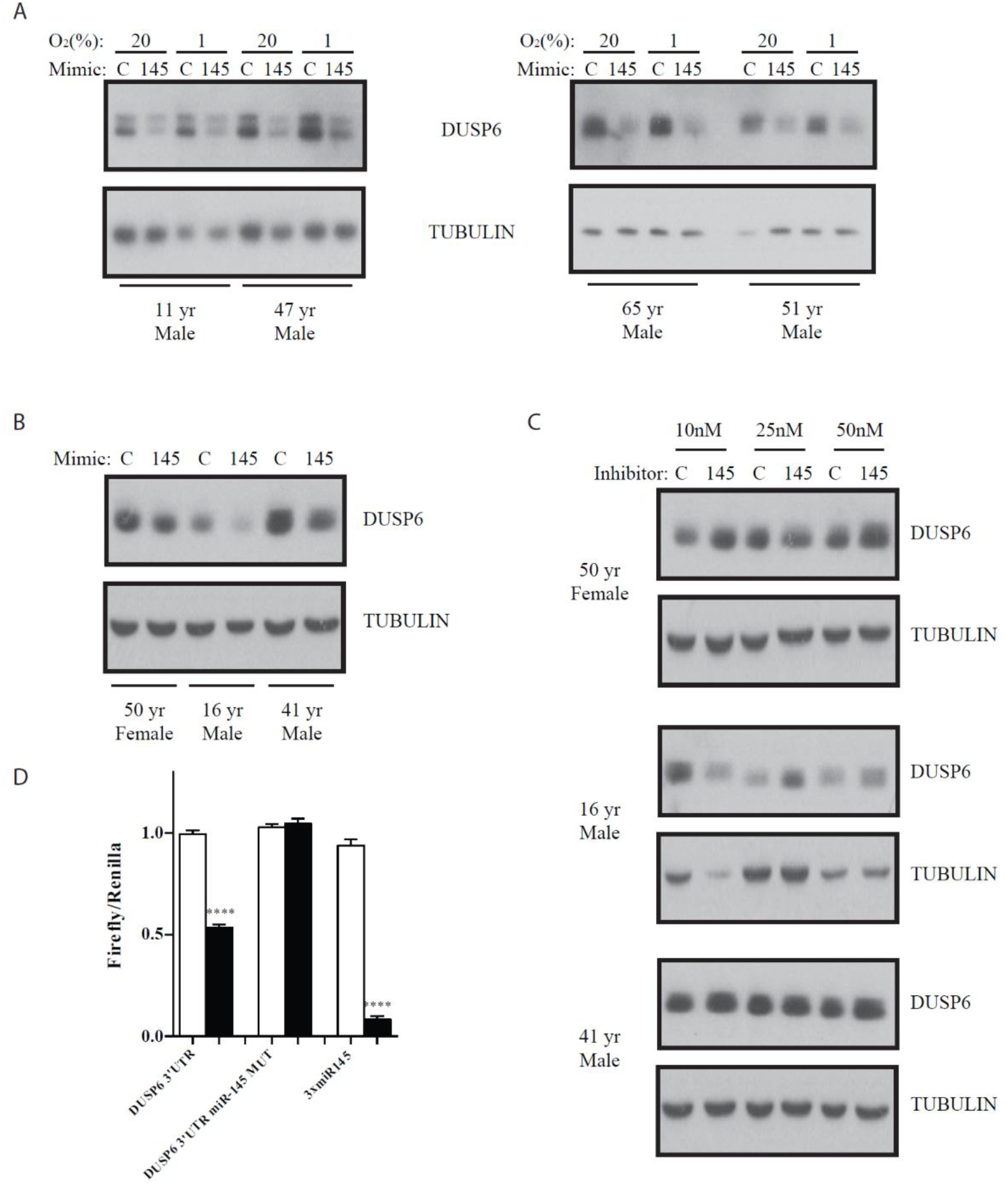
MiR-145 directly represses DUSP6. Western blot showing reduced DUSP6 levels following miR-145 over-expression in P2 **(A)** and freshly isolated **(B)** HACs. **C)** Western blot showing increased DUSP6 levels following miR-145 inhibition in freshly isolated HACs. In all experiments HACs were transfected with a relevant control (C) or miR-145 precursor or inhibitor, and subsequently cultured in 20% or, where indicated, 1% O2 tension for 44h. **D)** Hela cells were transfected with luciferase reporters containing three perfectly complementary binding sites for miR-145 (3xmiR-145), the DUSP6 3’UTR containing a putative miR-145 binding site (seed matching nucleotides 1018 to 1025; DUSP6 3’UTR) or a mutated seed-matching site (DUSP6 3’UTR miR-145 MUT) downstream the firefly luciferase open reading frame (ORF). Control (‘C’) or miR-145 mimics were co-transfected in the cells. Values were normalized to the levels of renilla luciferase, independently expressed by the same vector and are shown as relative to that obtained for each construct co-transfected with the control miRNA mimic (±S.E.M). \*\*\*\**P* < 0.0001.

**Table 2.**
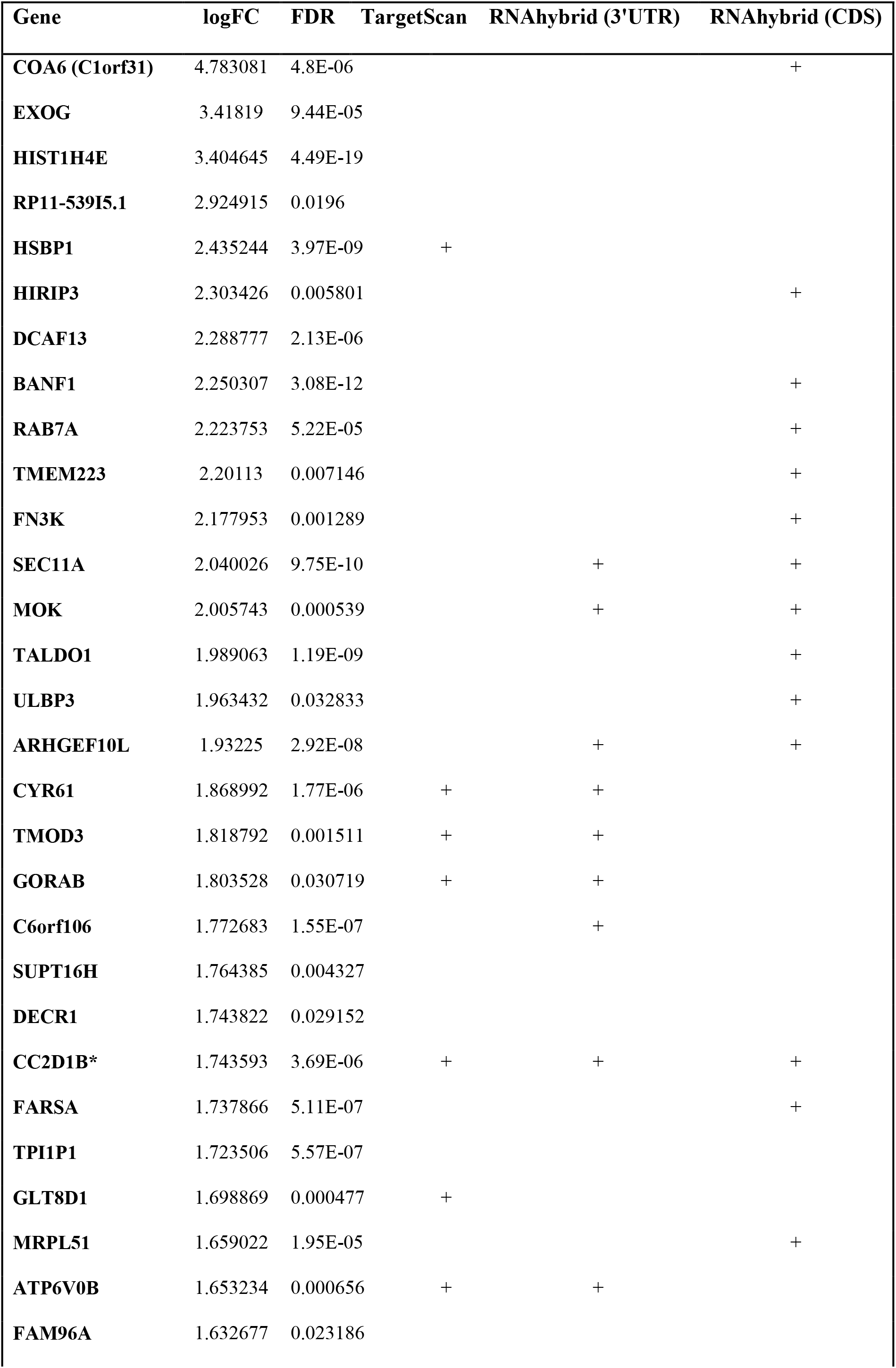
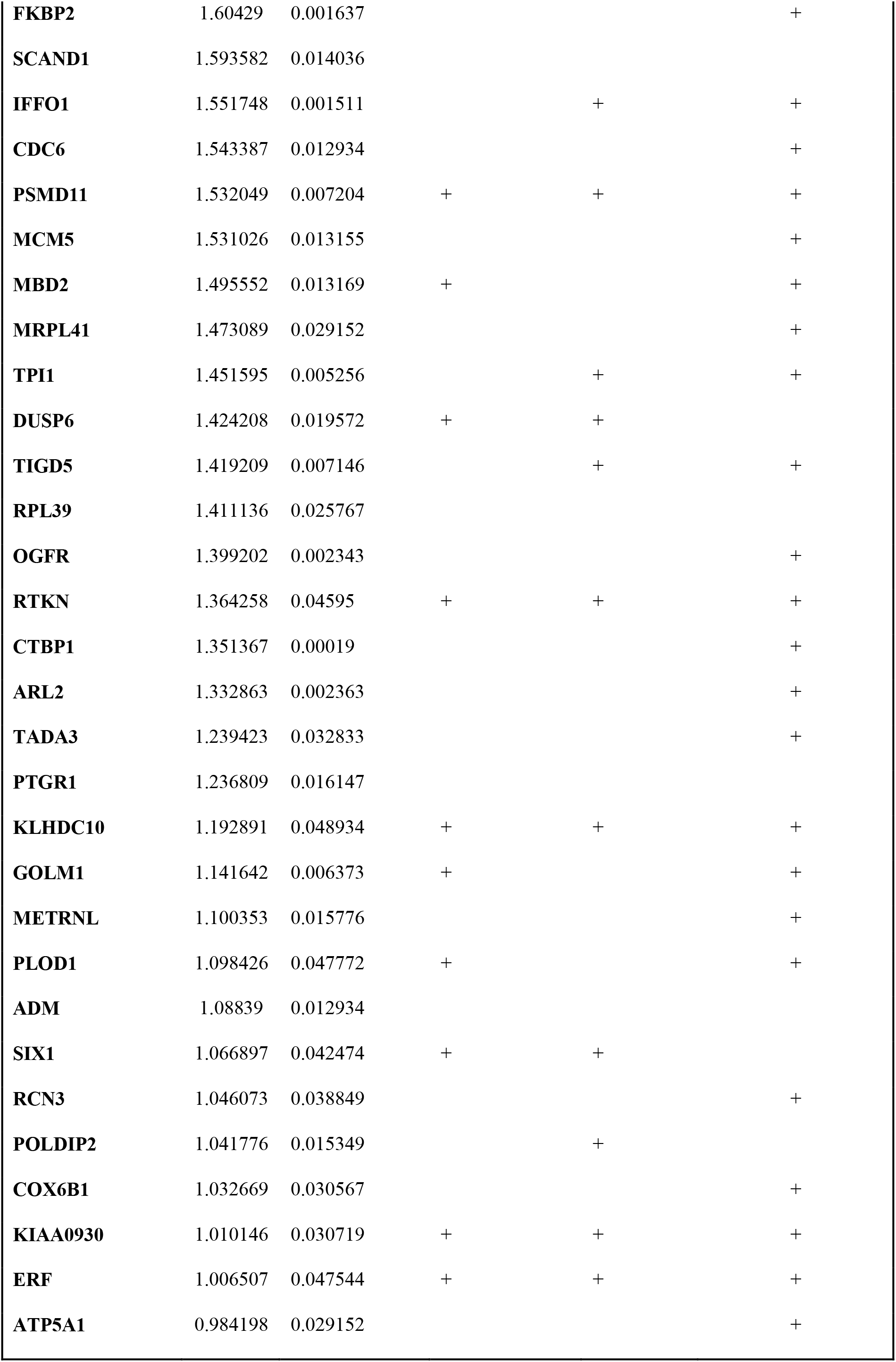
mRNAs significantly enriched (FDR<0.05) in Ago2-immunoprecipitates following miR-145 overexpression. (DT-IP: Direct targets (candidates)). Control or miR-145 mimics were transfected in P3 HACs from three different patients. 24h after transfection, cells were lysed and submitted to Immunoprecipitation with anti-Ago2 antibodies. Immunoprecipitated RNA was extracted and submitted to HT-seq. The list only includes those genes which expression was down-regulated or unchanged as a consequence of miR-145 overexpression. The last three columns indicate the presence of predicted binding sites (+) accordingly to TargetScan (3’UTRs) and RNAhybrid (3’UTRs or CDSs, as indicated).

FGF2 (Fibroblast growth factor 2) plays an anti-anabolic and/or catabolic role in HACS (31, 32) at least partially via the activation of the Ras-Raf-MEK1/2-ERK1/2 pathways(33). FGF2 is present in OA synovial fluid, where it activates Runx2 and promotes MMP-13 expression (2). Levels of FGF2 and MMP13 are increased in human OA cartilage and, in adult human articular cartilage, FGF2 stimulates MMP-13 through the activation of multiple MAPKKs. (34). Additionally, mice in which increased Erk1/2 phosphorilation occurred upon heparin endosulfatases (Sulf1 and Sulf2) depletion, showed increased MMP-13 levels and reduced Col2a1 and ACAN (35) expression. Interestingly, a raise in Runx2 and MMP13 and a reduction in Col2a1 and ACAN was observed upon miR-145 over-expression (11). In mouse embryos, transcription of DUSP6 is dependant of FGF signalling and targeted inactivation of DUPS6 increased the levels of pERK, the main target described for DUSP6. Thus, we hypothesized that DUSP6 could contribute to the effect of miR-145 in the expression of the catabolic/anabolic molecules mentioned above. Whereas DUSP6 depletion with specific siRNAs didn’t alter Col2a1 or ACAN levels (data not shown), expression of MMP-13 was strongly up-regulated (Figure 4A), indicating that miR-145-dependent MMP13 regulation may be partially mediated by DUSP6. Nevertheless, miR-145 overexpression or siRNA-mediated DUSP depletion failed to impair ERK phosphorylation in response to FGF2 (Figure 4B/C.

**Figure 4.**
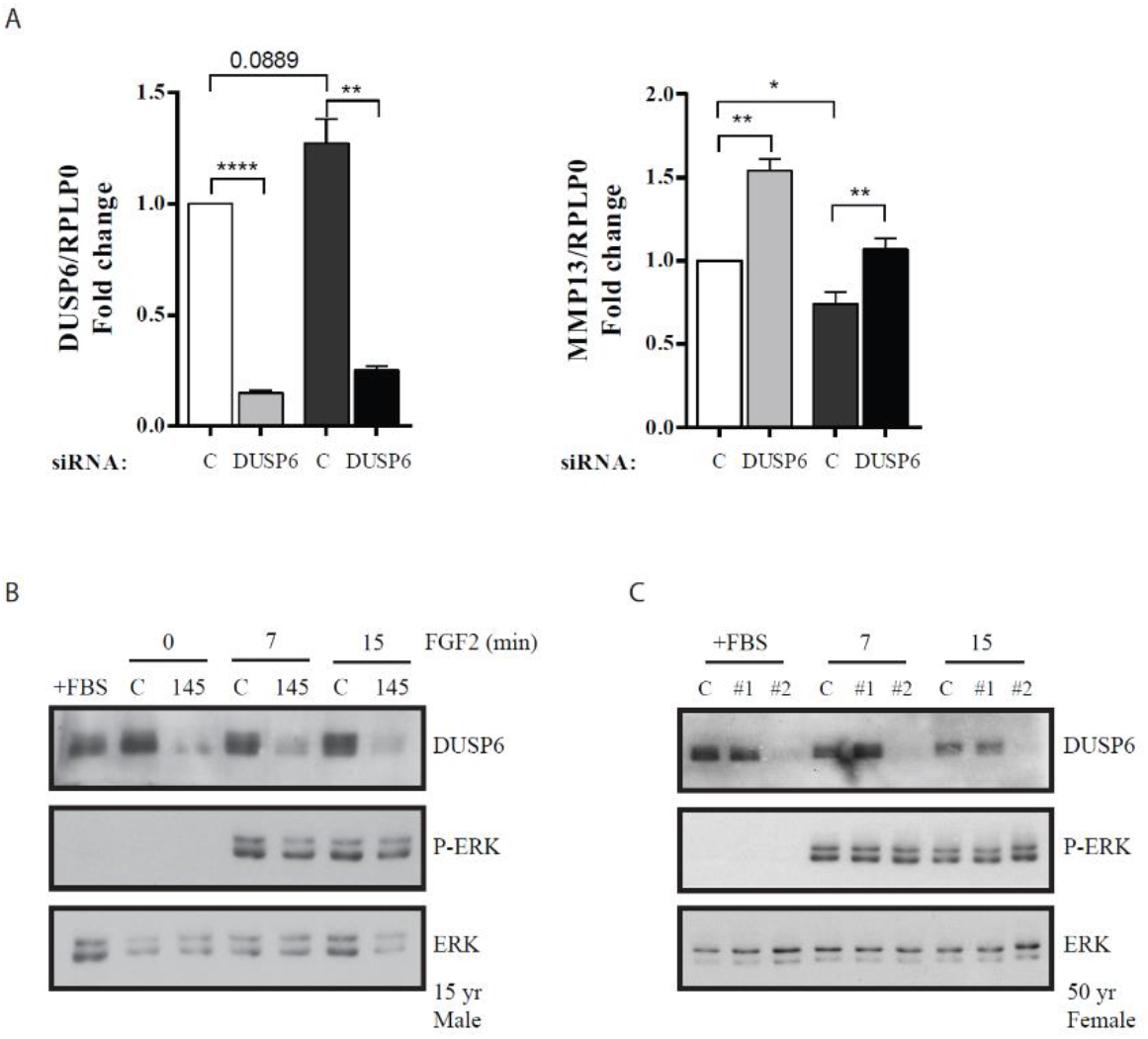
DUSP6 depletion results in MMP13 up-regulation. **A)** RT-qPCR analysis of DUSP6 and MMP13 levels after transfection of P2 HACs with 5nM DUSP6 siRNA. Values are presented as relative to that obtained in cells transfected with control siRNA for each patient and normalized to RPLP0. Data represents average ± S.E.M. from 4 different experiments, each performed with different donor cells. \**P* < 0.05; \*\**P* < 0.01; \*\*\*\**P* < 0.0001 **B, C)** Western blot showing ERK phosphorylation (pERK) after treatment with FGF2 (25nM). P1 HACs were transfected with control (C) or miR-145 (145) mimics **(B)**, or control (C) or DUSP6 siRNAs (#1, #2, notice than only #2 siRNA results in DUSP6 depletion,therefore #1 acts as an additional control) **(C)**. 24h later, HACs were placed in serum-free medium overnight and stimulated with 25nM FGF2 for the indicated times. These experiments were performed three times, with three different donors; one representative experiment is shown.

Altogether, these data indicate that miR-145 function can be modified without significantly interfering with FGF2-dependant ERK activation and the effect of DUSP6 in MMP13 expression occurs through a still un-identified ERK1/2-independent pathway.

## DISCUSSION

We have applied RIP-Seq in combination with total mRNA-Seq following miR-145 overexpression to successfully identify >50 new direct miR-145 targets in primary human articular chondrocytes. MiRNA-target interaction results in translational repression accompanied or not of mRNA degradation (36–38). Therefore, experimental approaches that rely on transcriptome profiling alone fail to detect targets regulated exclusively at the level of translation or to differentiate between direct and indirect targets. Surprisingly, most miR-145-mRNA interactions in HACs didn’t result in mRNA degradation 24h after overexpression. However, a reduction in several of these transcripts was observed at a later time point and luciferase assays confirmed that miR-145-target mRNA interaction results in lower protein (luciferase) levels for the majority of the cases (11 out of 13). Our results therefore suggest that miR-145 functions by inhibiting translation of most of its direct targets and is later followed by mRNA decay. The main mechanism by which miRNAs repress gene expression remains highly controversial (39) (36) (40). Interestingly, recent studies suggest that the mechanism of action might be cell-type specific (41) and dependent of the target site type. Thus, Liu and collaborators found that extensive complementarity to the seed region of the miRNA through the 3’UTR often results in mRNA decay instead translational repression (37). Nevertheless, it remains unclear which features favour mRNA translation over decay, (37). We demonstrate here that, in HACs, miR-145 binding occurs more often through CDSs. Recent studies have shown that interaction through the CDS occurs frequently (18, 26, 42). If the region of interaction miR-145/mRNA influences the mechanism of silencing in HACs is a possibility that remains to be clarified.

MiR-145 overexpression resulted in up-regulation of a high number of genes (272-Supplemental Table 1) and down-regulation of >80 genes, suggesting indirect effects, not entirely surprising, since, as we have shown in the past, miR-145 targets SOX9 (11), that regulates a wide range of genes in HACs. Further supporting this idea, only one of the down-regulated genes was enriched in Ago-IPs although several additional genes from this list were enriched in Ago-IPs upon exclusion of the outlier sample from the analysis. Thus, our immunoprecipitation protocol may fail to recover weak mRNA-miRISC interactions. mRNA crosslinking may increase the yield of recovered interactions, although could led to potential artifacts due to reduced cell lysis efficiency, introduce sequence biases, increase background or be incompletely reversible (43, 44). Increasing the number of IP experiments would definitely have resulted in a more comprehensive list of actual targets, but this was limited both by the availability of healthy primary chondrocytes and the high cost of the HT-Seq, rapidly decreasing as the technology is optimized. Importantly mRNA-RISC is a dynamic interaction which can be hugely affected by the cellular context: our experiments have been performed in primary human chondrocytes, instead in the more accessible cell lines or mouse models that are commonly used in miRNA studies in cartilage, increasing the reliability and applicability of our results.

Although it remains to be demonstrated, several of the newly identified miR-145 targets in HACs may play an important role in cartilage function. For example, POLDIP2 might modulate production of MMP-1 through NOX4 (45, 46). Also, PLOD1 might potentially be involved in collagen metabolism (47) and ERF, an ETS2 repressor factor, could indirectly modulate the expression of ECM components (48, 49). Interestingly, CYR61 has already been shown to play a role in chondrogenesis (50), although its function in adult cartilage remains to be clarified.

OA pathology is characterized by progressive cellular and molecular changes in joint tissues. While it has always been considered a degenerative joint disease influenced by mechanical stresses and ageing causing cartilage breakdown, discoveries in the recent years point to a critical contribution of inflammatory mechanisms (51). Chondrocytes in the joint can express and respond to cytokines and chemokines such as TNF-a, IL-1 or IL-6 and both IL-1β and Interferon-γ suppress COL2A1 transcription. Interestingly, Yang et al. have recently found that miR-145 is up-regulated in OA cartilage and chondrocytes and HACs treated with IL-1β contained higher miR-145 levels. Moreover, inhibition of miR-145 partially reversed the IL-Iβ-mediated COL2A1 and ACAN impairment as well as the stimulation of MMP-13(12). GO and KEGGS analysis of the genes affected by over-expression of miR-145 show enrichment in pathways involved in inflammatory and interferon responses. Not surprisingly, a role for miR-145 in modulating interferon response has already been suggested in very different cellular contexts such as bladder cancer (52) or fish INF-γ-induced immune response (53). Nevertheless, the mechanisms underlying the involvement of miR-145 in these pathways remain unknown. Whereas miR-145 induces INF-β by targeting the suppressor of cytokine signalling 7 (SOCS7), we didn’t detect an effect of miR-145 in SOCS7 expression or association with miRISC. Thus, miR-145 most probably modulates inflammatory response in HACs through a SOCS7-independet mechanism. We have previously proposed that miR-145 inhibition in OA cartilage results in increased synthesis of main ECM components such as COL2A1 and silencing of degradative enzymes through its target *Sox9.* Our new findings suggest that OA cartilage treated with miR-145 inhibitors would also benefit of a reduction in the inflammatory responses that contribute to cartilage degeneration. Additionally, we have demonstrated that DUSP6 is targeted by miR-145 and that the effect of this miRNA in MMP13 expression previously described by Yang and ourselves(11) might be at least partially mediated by DUSP6. In our hands, levels of pERK upon FGF2 stimulation were similar in the absence of DUSP6 and/or presence of miR-145, suggesting that DUSP6 effect on MMP13 expression is independent of FGF2-mediated ERK phosphorylation. Further studies, especially those mimicking cartilage diseases, are required to confirm these findings and to shed more light into the mechanism of action of this phosphatase in HACs.

MiR-145 is abundantly expressed in cartilage, but is also highly expressed in various mouse tissues such as aorta, where miR-145 affects its structure (54). Moreover, changes in miR-145 levels have been associated with various human diseases, including stroke (55), arteriosclerosis (56) or cancer (57, 58). Although miRNA-target interactions can be cell-specific, our list of targets in HACs could be of interest in other fields and some of the interactions may be relevant in other tissues or diseases. Supporting that idea, RTKN (Supplemental table 1) has been already proven to be targeted by miR-145 in breast cancer (57), FSNC1 (table 1) in colon cancer or melanoma (59, 60) and GOLM1 (Supplemental table 1) in prostate cancer (61).

In summary, we demonstrate that miR-145 directly targets dozens of genes in the human articular chondrocyte to strongly impact its function. Our results suggest that inhibition of miR-145 in osteoarthritic cartilage, alone or in combination with other therapies, could not only contribute to the restoration of the balance between catabolism and anabolism of the ECM, but might also limit inflammation of the joints. The effect of modulating the expression of miR-145 target genes for chondrocyte function will need to be carefully assessed prior to the development of therapeutic approaches targeting miR-145. Experiments performed *in vivo* will surely address the suitability of miR-145-based therapies for the treatment of OA in the near future.

## SUPPLEMENTARY DATA

Supplementary data available

## ACKNOWLEDGEMENTS AND FUNDING

We thank the High-Throughput Genomics Group at the Wellcome Trust Centre for Human Genetics (funded by Wellcome Trust grant reference 090532/Z/09/Z and MRC Hub grant G0900747 91070) for the generation of the Sequencing data.

